# *jouvence*, a new small nucleolar RNA required in the gut extends lifespan in Drosophila

**DOI:** 10.1101/441121

**Authors:** Lucille Mellottée, Stéphanie Soulé, Abdelkrim Arab, Chongjian Chen, Jean-René Martin

## Abstract

Longevity is influenced by various genetic and environmental factors, but the underlying mechanisms remain poorly understood. Here, we functionally characterise a new Drosophila small nucleolar RNA (snoRNA), named *jouvence* whose loss of function dramatically reduces lifespan. A transgene containing the genomic region of *jouvence* rescues the longevity in mutant, while its overexpression in wild-type flies increases lifespan. *Jouvence* is expressed in epithelial cells of the gut. Targeted expression of *jouvence* specifically in the enterocytes increases lifespan, indicating that its role in the control of longevity takes place in these cells. A transcriptomic analysis performed from the gut reveals that several genes are either up-or down-regulated in mutant indicating that the snoRNA-*jouvence* might be involved in transcriptional control. Finally, since snoRNAs are structurally and functionally well conserved throughout evolution, we identified putative *jouvence* orthologue in mammals including humans, suggesting that its function in longevity might be conserved through evolution.

---

The aging process is the result of complex biological mechanisms leading to accumulation of different types of damage at molecular, cellular, tissue and organ levels. This leads to the decrease or loss of certain physiological functions, and therefore, to an increase of vulnerability to diseases and death [1,2]. Several genes and metabolic factors have been reported to modulate longevity [3–20]. Indeed, genes that increase longevity have been identified in different model organisms such as yeast [3,4), *C. elegans* [21], *Drosophila* [5–20] and mouse [22]. More particularly, in *Drosophila,* these include *mathuselah* [5], insulin signalling pathway genes [6–9,18,19], Cu/Zn superdismutase [10], sirtuin [3,4,11], and genes affecting either the regulation of mitochondria as PGC-1 [12] or the mitochondrial respiratory chain [13]. It is also well accepted that lifespan is environmentally modulated. In addition, longevity positively correlates with the ability to resist to stress. In such way, environmental factors like diet [1,15], food-derived odours [16] and stress [5,8,16,17] also modulate an organism’s lifespan. More recently, some microRNAs (miRNAs) were identified as key regulators of genes involved among others processes as neurodegeneration, in aging [23–26].

The snoRNAs are members of a group of non-coding RNAs. SnoRNAs, found in species from archeobacteria to mammals [27], are known to be present in the nucleolus, in which they are associated with a set of proteins to form small nucleolar RiboNucleoProtein (snoRNPs) [27-28]. They are generally known to be processed from introns of pre-mRNAs and are believed to have several functions, including the 2’-O-methylation and pseudouridylation of different classes of RNA, the nucleolytic process of the rRNA, and telomeric DNA synthesis [27–30]. Based on conserved secondary structure and functional RNA motifs, two major classes have been distinguished, the box C/D and box H/ACA. The box C/D, whose are the most known, comprising notably the multiple “U” snoRNAs, performs the 2’-O-methylation of ribosomal RNA. The less-known subtype, the box H/ACA directs the conversion of uridine to pseudouridine, and so particularly of rRNA [27]. However, recent studies have demonstrated that the H/ACA snoRNA might also pseudouridinylate other RNA substrates, such as mRNA and long-non-coding-RNAs (lncRNAs) (31) or have a role in chromatin remodeling, indicating that they have several other functions [32]. Then, although few H/ACA box snoRNA have been involved in human diseases [33,34], their role at the organismal level and more particularly in aging has not yet been documented.

Here, we have characterized a new small nucleolar RNA (snoRNA:Ψ28S-1153) that we have named *jouvence (jou)* and showed that its mutation reduces lifespan. A transgene containing the genomic region of *jou* is able to rescue the lifespan, while its overexpression increases it. *In-situ* hybridization (ISH) revealed that *jou* is expressed in the enterocytes, the main cell-types of the epithelium of the gut. Thus, genetic targeted expression of *jou* in these cells is sufficient to rescue and significantly increase longevity, and even only in adulthood (through Gene-Switch conditional expression). Finally, as snoRNAs are generally well conserved throughout evolution, both structurally and functionally, we have identified putative *jou* mammalian homologues, both in mouse and human, suggesting that it may have an implication in mammalian aging.

## RESULTS

### Genetic and molecular characterization of a new snoRNA

Aging involves a progressive decline and alteration in tissues and sensory-motor functions [35–37]. In a study to characterize putative genes involved in aging and particularly in aging effects on sensory-motor performance, a P-element insertional mutagenesis was performed and identified the enhancer-trap line P[GAL4]4C. P[GAL4]4C is inserted on the second chromosome, at position 50B1 (Figure 1A), between two new putative genes: CG13333 and CG13334. CG13333 is embryonically expressed [38], while CG13334 shows a partial and weak homology to the lactate dehydrogenase gene [39]. Furthermore, a bioinformatic study [40] has annotated a novel snoRNA (snoRNA:Ψ28S-1153) between CG13333 and CG13334 near the P[GAL4]4C insertion (Figure 1A), while a recent developmental transcriptomic analysis has annotated two additional putative snoRNAs (snoRNA:2R:9445205 and snoRNA:2R:9445410) (http://flybase.org/) [41] localized just upstream to the first snoRNA:Ψ28S-1153. However, according to the canonical definition of snoRNA in which the H/ACA box is predicted to form a hairpin-hinge-hairpin-tail structure (Figure 1D), while the C/D box is predicted to form a single hairpin [27-30], these two last snoRNAs do not fulfil these two criteria, suggesting that they are likely not true or, at least canonical snoRNAs. Whereas the snoRNA:Ψ28S-1153 forms a typical H/ACA double hairpin (Figure 1E). To generate mutations in this complex locus, the P[GAL4]4C was excised, using the standard genetic method of excision [42] resulting in a small 632bp deletion (named F4) Figure 1A and S1A), which completely removed the snoRNA:Ψ28S-1153 as well as the other two RNAs. Next, we performed RT-PCR to test if this deletion had an effect on the expression of the two encoding neighbouring genes. Both genes were normally expressed in the deletion line compared to control Wild-Type (WT) Canton-S (CS) flies (Figure S1B).

**Figure 1.**
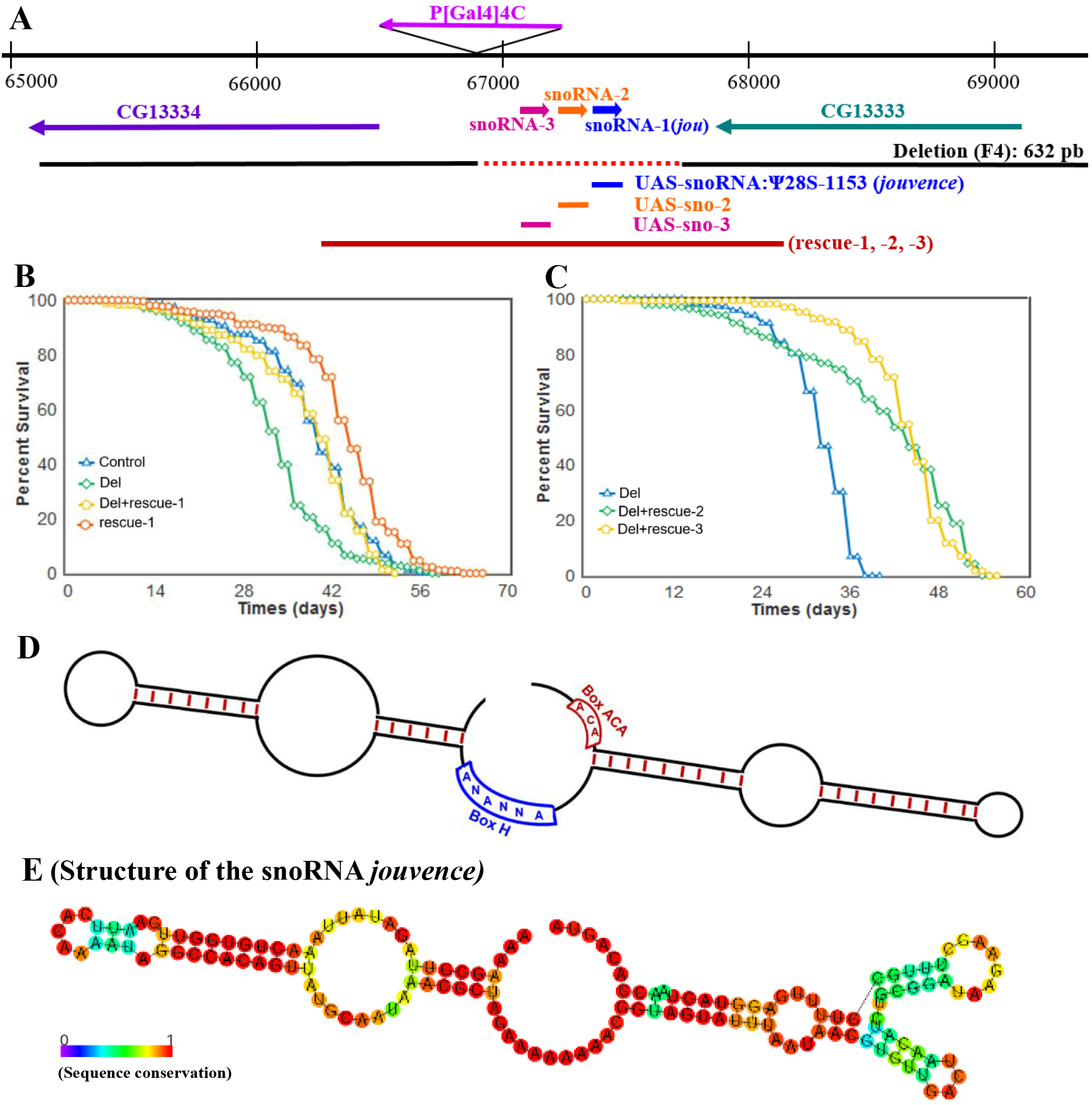
Molecular map of the P[Gal4] locus and features of the snoRNA:Ψ28S-1153. (A) Genomic map of the P[Gal4]4C locus. The snoRNA:Ψ28S-1153 (*jouvence*) as well as the two other putative upstream snoRNAs (sno-2=snoRNA:2R:9445205 and sno-3= snoRNA:2R:9445410) are in the inverse orientation of the two encoding genes: CG13333 and CG13334. The deletion (F4) of 632 bp (red dotted line) including the three putative snoRNAs. The 1723 bp genomic DNA fragment (genomic-rescued named: rescue-1, rescue-2 and rescue-3) (red line) used to generate transgenic flies, the snoRNA *jou* fragment (blue bar) of 148bp used to generate the UAS-jou construct, the 166bp fragment (orange bar) corresponding to the sno-2, and the 157bp fragment (pink bar) corresponding to the sno-3. B) Decreasing cumulative of Control (CS), Del (deletion F4), Del+rescue-1 and rescue-1. Note: the lifespan determination of these 4 groups of flies (genotypes) have been performed simultaneously, in parallel (for Statistics: see Table-S1). C) Similarly to B, decreasing cumulative of females Del (deletion F4), Del+rescue-2, Del+rescue-3. In Del+rescue-2, Del+rescue-3, the lifespan is lengthened (for Statistics: see Table-S1). D) Schematic representation of a H/ACA snoRNAs structure. E) Schematic representation of the snoRNA:Ψ28S-1153 (*jouvence*), harboring a typical H/ACA box structure.

### Deletion of the genomic region encompassing snoRNA:Ψ28S-1153 reduces lifespan

Interestingly, we observed by daily husbandry, that the deletion flies had a shorter lifespan. Hence we compared the longevity of these flies to that of Control CS. Since small variations in the genomic background can affect longevity [8,16], we outcrossed the deletion line a minimum of 6 times to CS (Cantonisation). Longevity tests (Figure 1B) revealed that female deletion flies lived shorter compared to control. To test the snoRNA’s role in the longevity effect, three transgenic lines containing a DNA-fragment of 1723 bp, comprising the putative genomic and regulatory region of the snoRNAs (genomic-rescued named: rescue-1, 2 & 3) (Figure 1A: red bar) were generated, and tested for longevity against the deletion background. Again here, these transgenic lines were outcrossed 6 times to CS and then crossed to the deletion to generated the Del+rescue-1 line. Longevity in Del+rescue-1 females was rescued, while lifespan in the rescue-1 line itself increased (Figure 1B). Similar results were observed with the two other independent insertions: rescued-2 and rescue-3 (Figure 1C). In addition, as aforementioned, since longevity is very sensitive to the genetic background, we also tested the phenotype of the deletion in an independent genetic background (Berlin) [43].

The deletion flies in a Berlin genetic background still live shorter than their co-isogenic Berlin flies (Figure S1D), corroborating that deletion of this genomic region reduced lifespan and was not due to a bias in the genetic background.

### The snoRNA:Ψ28S-1153 is expressed in the epithelium of the gut

Since only the snoRNA:Ψ28S-1153 has been identified in the bioinformatic study as a real/canonical snoRNA [40], we hypothesize that this snoRNA might be potentially the only true functional snoRNA in this locus. In order to determine in which tissues snoRNA:Ψ28S-1153 could be expressed and required to affect lifespan, we performed *in-situ* hybridization (ISH) on the whole flies using fluorescently-labelled probes (Figure 2). We then performed ISH with this probe and found that it was expressed in gut epithelial nucleoli and the proventriculus (Figure 2A), but as expected, its expression was absent in deletion line (Figure 2A2). Moreover, a counter-staining with DAPI suggest that it is localised in the nucleolus (Figure 2A3-4). In addition, whole body examination revealed that the snoRNA:Ψ28S-1153 was also expressed in ovarian nurse cell nucleoli in control flies (Figure S2), but not in any other tissues, including the nervous system (see Figure S3A for the expression pattern of the whole fly, including the head, the nervous system and the thorax). Similarly, when the rescue construct was combined with the deletion line, snoRNA:Ψ28S-1153 expression was restricted to gut epithelial cells, though surprisingly, not in the ovaries. Additional ISH performed on dissected gut of Wild-Type flies confirmed that the snoRNA:Ψ28S-1153 was expressed more precisely in the midgut (Figure 2C,D), but not in the hindgut (Figure 2E). These results suggested that effects on longevity of snoRNA:Ψ28S-1153 were likely due to its expression in the gut epithelial cells, and not in the ovaries, although at this stage of this study, the role of the two other putative snoRNAs could not be excluded.

**Figure 2.**
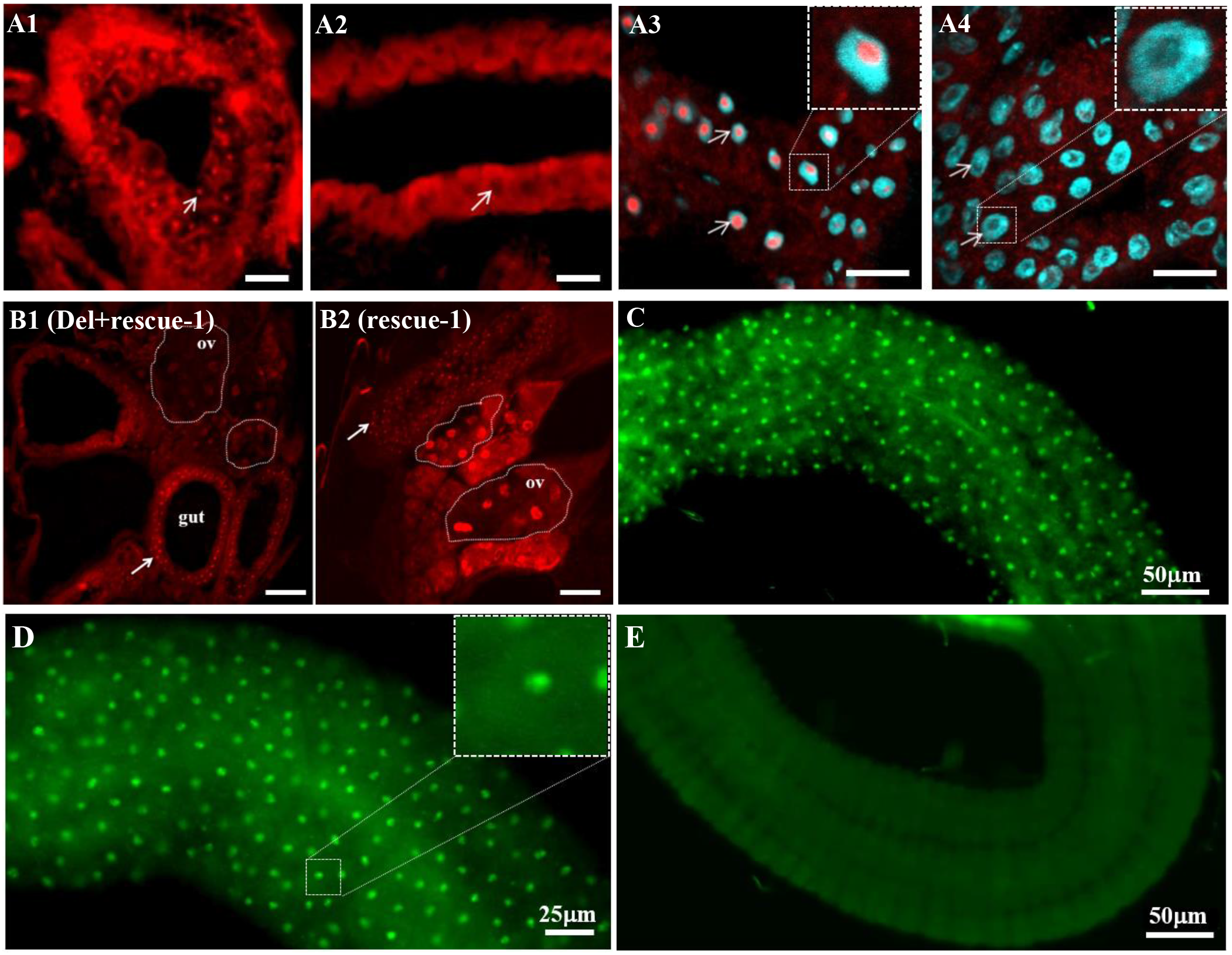
snoRNA:Ψ28S-1153 *(jou)* is expressed in the epithelium of the gut. (A) *in situ* Hybridization (ISH) of the snoRNA:Ψ28S-1153 on whole fly (Cryosat section) reveals a restricted expression in the epithelium of the gut in Control (A1), but it is absent in the deletion (A2). In Control, the snoRNA was detected with a granular expression pattern (red dots: white arrow), but in deletion, no expression. (A3) and (A4) Magnified view taken with a confocal (63X) revealed that *jou* is expressed in the nucleolus (DAPI counter-staining), but absent in deletion (A4) (scale bar = 25μm). Inset (magnification) in A3 and A4 showing that *jou* is located in the nucleolus. (B) ISH (cryostat section) of *jou* on transgenic whole flies (Del+rescue-1 and rescue-1). In rescue-1 (Wild-Type background) (B2), *jou* expression was seen in the gut epithelium (red dots: white arrow) and ovaries (ov) as expected (see Figure S2), while expression was restricted to the gut epithelium in Del+rescue-1 and absent in the ovaries (ov) (scale bar = 50μm). C,D,E) ISH on dissected gut of Wild-Type CS flies. The ISH are revealed with a FITC-labelled tyramide (green). In C,D, which correspond to the midgut, the snoRNA is perfectly detectable in each cells of the epithelium. Inset (magnification) in D shows that *jou* is visible in the nucleolus even without specific labelling. In E, which corresponds to the hindgut, the snoRNA *jouvence* is not expressed.

The gut epithelium is composed of four cell-types: enterocytes (ECs), enteroblasts (EBs), entero-endocrine cells (EEs) and intestinal stems cells (ISCs) [44–49]. We specifically marked these cell types by combing *in-situ* Hybridisation (ISH) expression with GFP expression driven by cell type-specific drivers: ECs marked by Myo1A-Gal4, EBs detected through Su(H)GBE-Gal4 [48,49]), and ISCs marked by both esg-Gal4 and Dl-Gal4. Amongst these, only EC-specific expression (Myo1A-Gal4 showed *jou* co-labelling with the GFP reporter (Figure S4), indicating that its localisation was restricted to enterocytes.

### The snoRNA:Ψ28S-1153 in adulthood rescues and extends lifespan

To corroborate the previous results obtained with the genomic rescued transgene (Del+rescue-1) and confirm that the longevity effect is due to the snoRNA-Ψ28S-1153 in the epithelial cells of the gut, and not to the two others putative snoRNAs, we generated a p[UAS-snoRNA-Ψ28S-1153] transgenic line. Then, we targeted this unique snoRNA expression specifically to enterocytes in deletion background using Myo1A-Gal4 line (Del, Myo1A>UAS-snoRNA-Ψ28S-1153) and verified by ISH that the snoRNA was indeed correctly expressed in the enterocytes (Figure S4E). Furthermore, the expression of the snoRNA-Ψ28S-1153 specifically in enterocytes was sufficient to increase the longevity of the flies (Figure 3A), compared to their two respective controls (Gal4 line alone in heterozygous and UAS-jou alone in heterozygous). In addition, to investigate if the two other putative snoRNAs (snoRNA:2R:9445205 and snoRNA:2R:9445410, short-named sno-2 and sno-3 respectively) localized just upstream to the snoRNA:Ψ28S-1153 were also involved in the lifespan determination, the targeted expression of snoRNA-2 and snoRNA-3 in ECs did not increase lifespan of the deletion line encompassing snoRNA:Ψ28S-1153, indicating that only the snoRNA:Ψ28S-1153 expression in ECs is able to rescue lifespan among these three snoRNAs. To further support this demonstration, we also found a rescue of longevity defects compared to their co-isogenic control flies by expression of snoRNA-Ψ28S-1153 with a second Gal4 driver line, Mex-Gal4 [50], which is also expressed in the epithelium of the gut though not exclusively in the ECs [51] (Figure 3D). In the deletion background, the targeted expression in the epithelial cells of the gut by Mex-Gal4 also increases lifespan.

**Figure 3.**
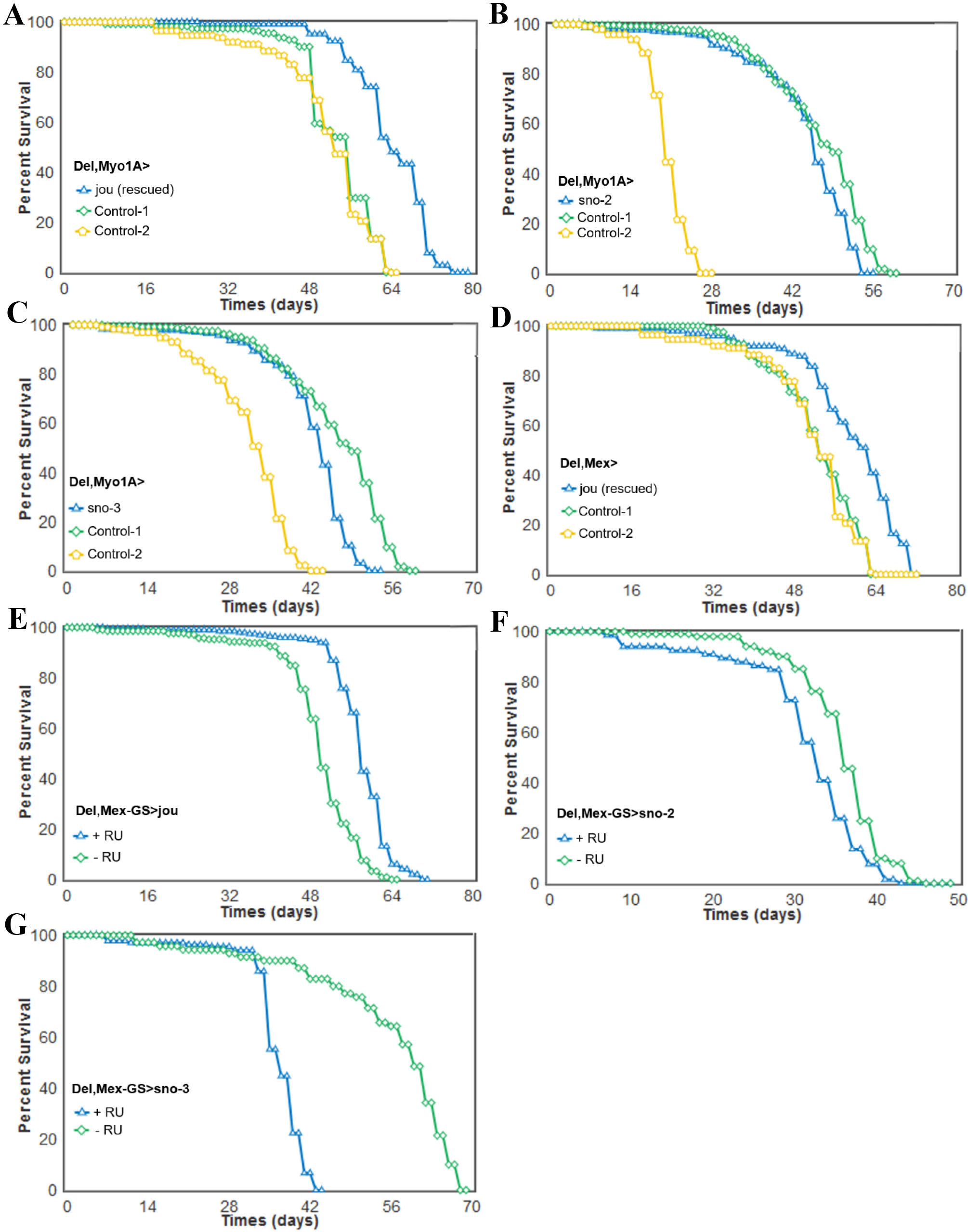
Targeted expression of the snoRNA:Ψ28S-1153 in the enterocytes is sufficient to rescue the longevity in deletion. Longevity test results (survival curve-decreasing cumulative) of the targeted expression of the snoRNA:Ψ28S-1153 specifically in the enterocytes in deletion compared to their controls. A) Del,Myo1A-Gal4>UAS-jou expression in enterocytes is sufficient to increase lifespan. B,C) The expression of the two other snoRNAs: sno-2 (B), or the sno-3 (C), does not lead to an increase of lifespan, and even it seems to be slightly deleterious. D) A second gut driver line, Mex (Del,Mex-Gal4>UAS-jou) which also targets the expression in enterocytes is sufficient to increase lifespan. E) The conditional Mex-GS flies (Del,Mex-GS>UAS-jou) fed with the RU486 only in adulthood, which triggers the expression of the snoRNA-*jou*, is sufficient to increase lifespan compared to the non-fed (non-induced) sibling flies. F,G) Mex-GS flies fed with RU486 only in adulthood, which triggers the expression of the snoRNA-2 (F) or snoRNA-3 (G), do not increase lifespan compared to the non-fed (non-induced) sibling flies, but it is rather deleterious (for Statistics: see Table-S1).

We then wanted to exclude that expression of snoRNA:Ψ28S-1153 by Myo1A-Gal4 and the Mex-Gal4 during development rescues longevity defects. We therefore generated a conditional Gal4 Gene-Switch line [52,53] (Mex-GS) to allow expression only in adult stages. In the deletion genetic background, feeding flies with RU486 starting just after the hatching and so, during adulthood, led to an increase of lifespan compared to the sibling control flies without RU486 (Figure 3E). In contrast, similar RU486 inducible experiments targeting either the UAS-sno-2 or the UAS-sno-3 did not yield to an increase of lifespan but rather to a decrease of lifespan (Figures 3F,G), indicating that the targeted expression of these two snoRNAs respectively could be deleterious, and consequently confirming that *jouvence* is the snoRNA responsible for the lengthening of lifespan.

To correlate snoRNA:Ψ28S-1153 expression levels and longevity, we performed a quantitative PCR on the dissected gut (Figure 4A). As expected, the snoRNA:Ψ28S-1153 is not detected in deletion line, whereas its expression is rescued in genomic-rescued line (Del+rescue-1) compared to Control. Moreover, its expression is significantly increased (4 folds) in rescue-1 transgenic flies, confirming overexpression in this line that carries four copies of the snoRNA-*jouvence*. We also quantify the level of the snoRNA on dissected gut (Figure 4B) in the two gut-driver lines. The levels of the snoRNA:Ψ28S-1153 was increased about 500-fold for the Myo-1A and about 600-fold with Mex-Gal4. Similarly the Mex-Gene-Switch driving the snoRNA-*jou* (Del,Mex-GS>UAS-jou) induced an increase of *jouvence* after RU486 induction (up to 5 folds) (Figure 4C). Thus, increased levels of snoRNA:Ψ28S-1153 were always correlated with increased longevity.

**Figure 4.**
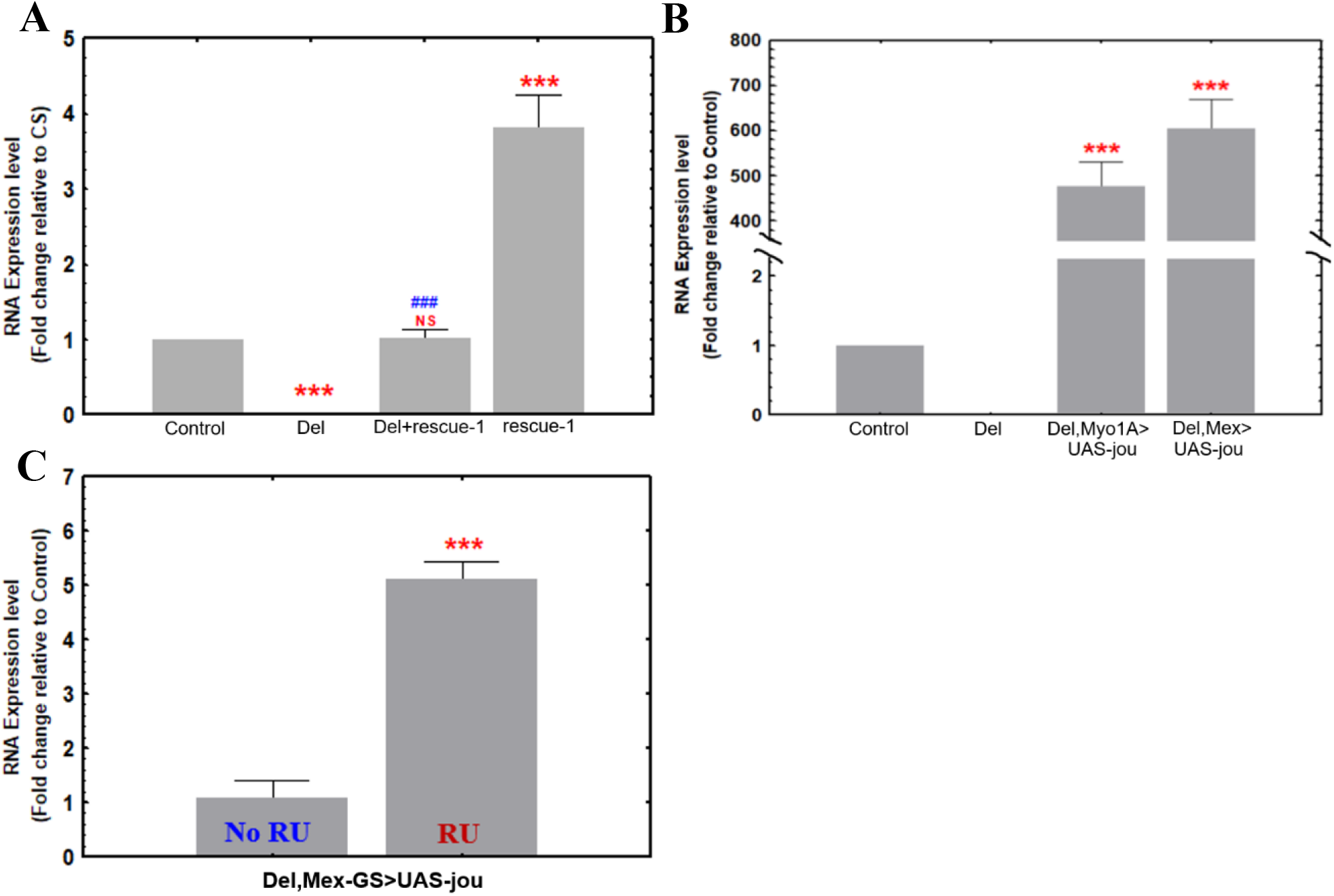
Histogram of RT-qPCR (Taqman) showing the relative levels of *jou* expression from the total-RNA of the gut. A) Control (CS), Del (deletion F4), genomic-rescued in the deletion (Del+rescue-1), and genomic-rescued transgene in the Wild-Type (rescue-1) (overexpression) (normalised and compared to Control-CS = 1). B) Quantification of the targeted expression of *jou* specifically in the enterocytes, both in Myo1A-Gal4 and Mex-Gal4 driving the UAS-*jou* in the deletion background, compared to the deletion. The expression is highly increased in flies expressing *jou* (Myo1A-Gal4 and Mex-Gal4). C) The conditional line Mex-GS flies fed with RU486 only in adulthood confirms the induction of the expression of *jou* compared to their sibling flies without RU486. Statistics: RU= Relative Units. Statistics: (p-values) (* compared to CS, # compared to F4). * or # p<0,05; ** or ## p<0,005; *** or ### p<0,0005).

### Overexpression of the snoRNA:Ψ28S-1153 in enterocytes of WT flies increases lifespan

Since in the deletion background, the targeted expression of the snoRNA:Ψ28S-1153 specifically in the enterocytes rescued the phenotype, we wondered if the overexpression of this snoRNA in a Wild-Type genetic background (CS) could be beneficial leading to an increase of lifespan or in contrast, be deleterious. We used the same driver lines to overexpress the snoRNA:Ψ28S-1153 up to 13-fold (Myo1A-Gal4) and 28-fold (Mex-Gal4) over that of endogenous expression (Figure 5F). Both Gal4-drivers expressing the snoRNA gave similar results, showing increased longevity (Figure 5A,B), compared to their co-isogenic controls flies. Moreover, conditional adult-specific expression via RU486 feeding in the Mex-GS line (30X increased expression) also increased lifespan compared to the noninduced flies (Figure 5C, 5G). Finally, the induction of the overexpression in adulthood, by the Mex-GS, of the sno-2 or the sno-3 did not alter lifespan compared to their respective co-isogenic control flies (Figure 5D,E), corroborating that the snoRNA-jou is critical for lifespan determination. In conclusion, since the snoRNA:Ψ28S-1153 increases lifespan when it is overexpressed in the epithelium of the gut, we conclude that it could be considered as a longevity gene, and therefore we named it “*jouvence*” (*jou*) (which means “youth” in French).

**Figure 5.**
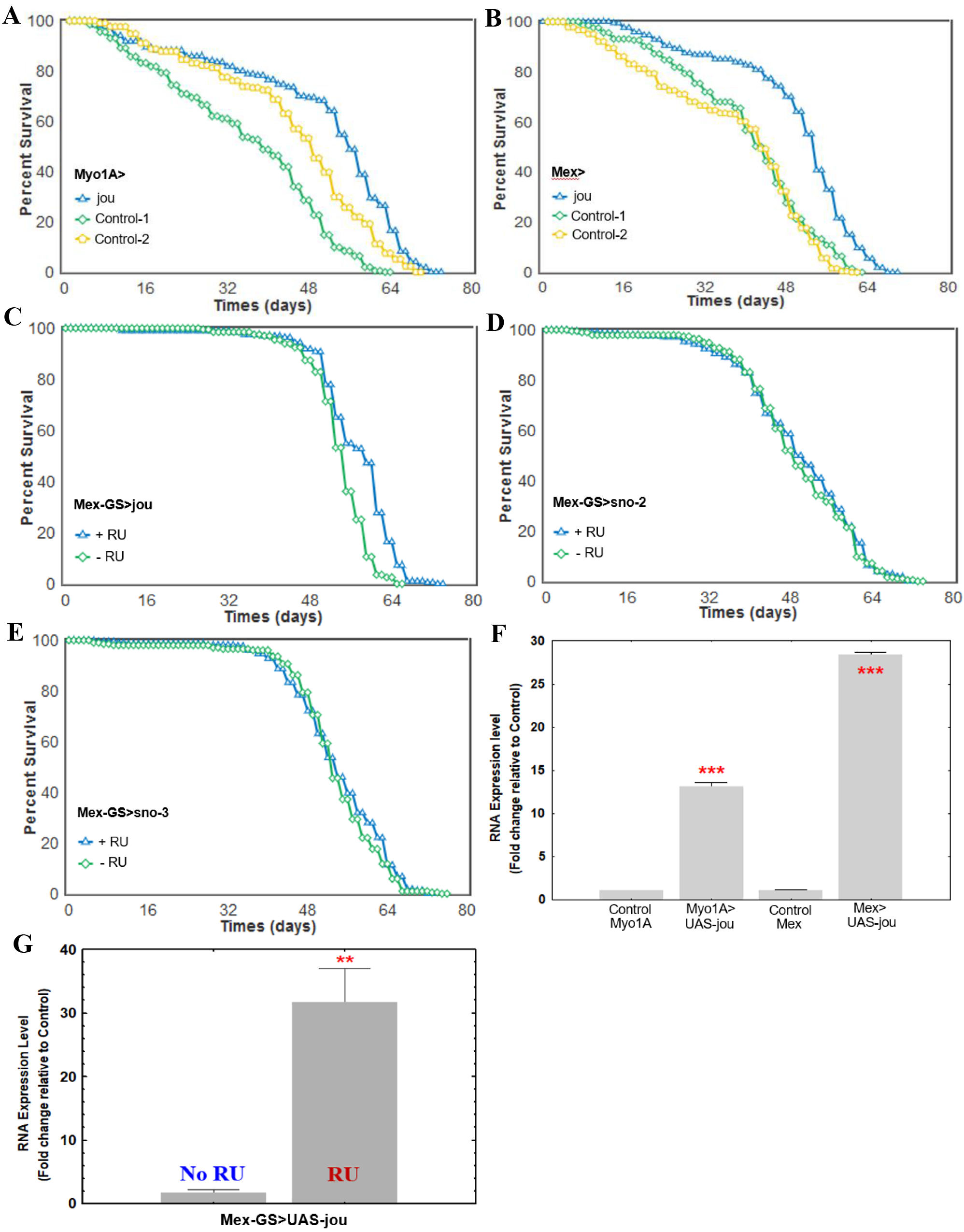
Overexpression of *jou* in the enterocytes increases lifespan. Longevity test results (survival curve-decreasing cumulative) of the targeted expression of *jou* specifically in the enterocytes in Wild-Type genetic background compared to their controls. Myo1A-Gal4>UAS-*jou* (A) and Mex-Gal4>UAS-*jou* (B) overexpression in enterocytes is sufficient to increase lifespan. C) Overexpression of *jou* only in adulthood is sufficient to increase lifespan (Mex-GS>UAS-*jou* flies fed with RU486 only in adulthood). D&E) Overexpression of the sno-2 (D) or sno-3 (E) only in adulthood do not have any effect on lifespan (Mex-GS>UAS-sno-2 or sno-3 flies fed with RU486 only in adulthood) (for Statistics: see Table-S1). F) RT-qPCR (Taqman) results of the targeted overexpression of *jou*specifically in the enterocytes compared to their controls Myo1A-Gal4, and Mex-Gal4, respectively. G) RT-qPCR (Taqman) results of the RU486 induced targeted overexpression of *jou* specifically in the enterocytes only in adulthood in Mex-GS, compared to the non-induced controls flies. Statistics: compared to control Myo/CS, or Mex/CS, or to the non-induced Mex-GS: (p-values) (* p<0,05; ** p<0,005; *** p<0,0005).

### The snoRNA *jou* is conserved through evolution

The snoRNAs are well conserved, both structurally and functionally, throughout evolution from archeabacteria to humans [27–29]. Bioinformatic analyses using BLAST and tertiary structure motif searches (using the Infernal software) revealed *jou* homologues in 12 *Drosophila* species, two orthologs in mice and one in human (Figure 6), suggesting positive evolutionary pressure for this particular snoRNA. In mouse, putative *jouvence* snoRNAs are located on chromosome 15, at position: 30336889-30337017, and on chromosome 18 at position: 6495012-6495133, respectively. The unique putative snoRNA *jouvence* in human is located on chromosome 11, at position: 12822722-12822880. Moreover, in Drosophila species, the two other putative snoRNAs (snoRNA-2 and snoRNA-3), which are highly homologous (more than 90%) (Figure S6) were identified just upstream to the snoRNA *jouvence* in 11 over the 12 other Drosophila species, except in *Drosophila grimshawi* (the most divergent from *D. melanogaster*), in which only the snoRNA-2 is present. However, these two putative snoRNAs have not been identified upstream to the *jou* homologue in mammals, suggesting that these two snoRNAs are likely not directly functionally linked to *jouvence*.

**Figure 6.**
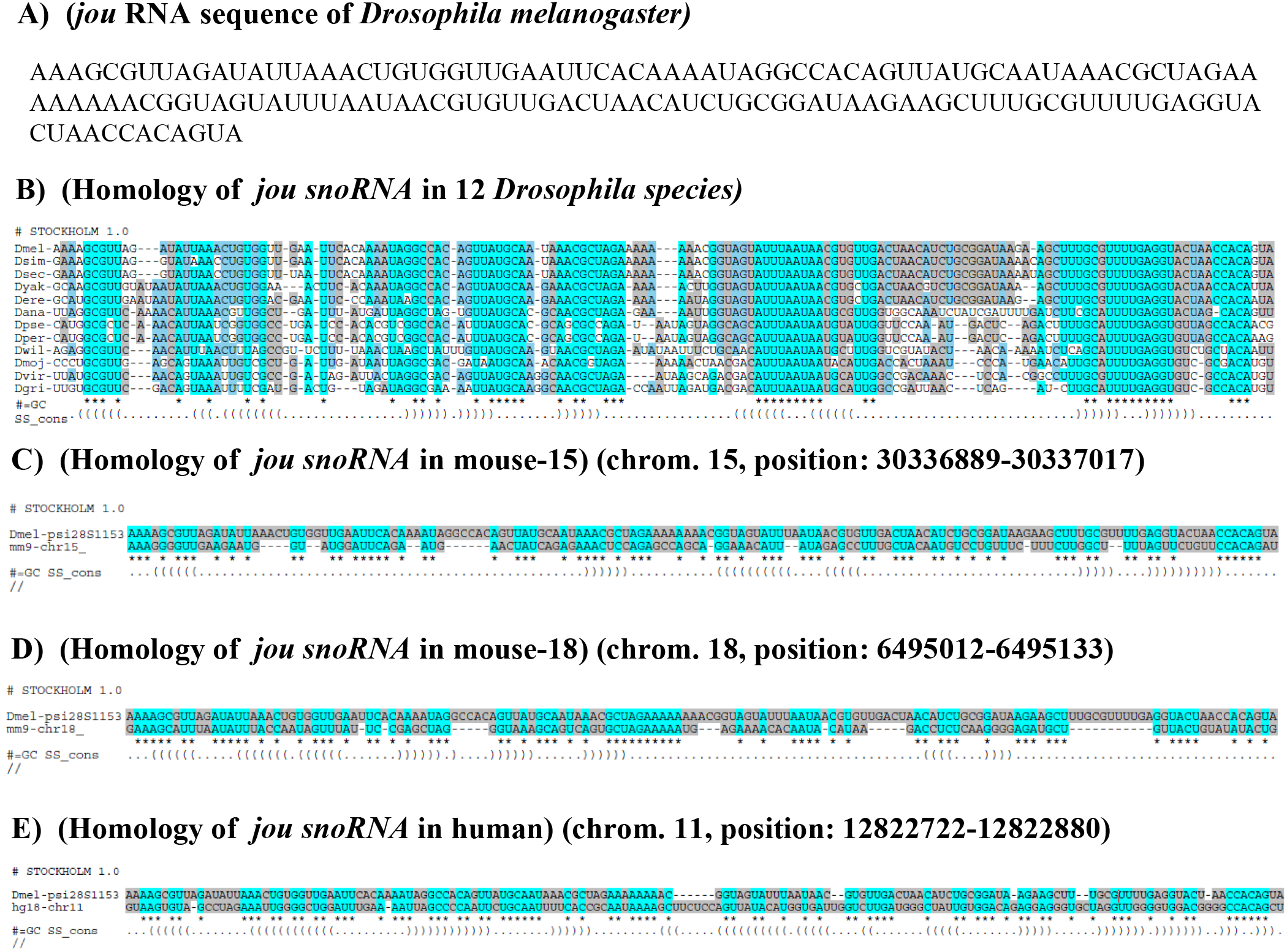
*jou* is conserved throughout evolution. A) RNA sequence of the *Drosophila jou* snoRNA. B) Homology of *jou* sequences in 12 *Drosophila* species available on FlyBase. Dmel= *Drosophila melanogaster*, sim = *simulans*, sec = *sechellia*, yak = *yakuba*, ere = *erecta*, ana = *ananassae*, pse = *pseudoobscura*, per = *persimilis*, wil = *willistoni*, moj = *mojavensis*, vir = *virilis, gri = grimshawi.* C&D) Ortholog sequences of *jou* in the mouse genome (*Mus musculus*). Two *jou* copies were detected, located on chromosomes 15 and 18. E) Ortholog sequence of *jou* in the human genome. Only one copy was identified, located on chromosome 11.

To explore further a putative role of the snoRNA *jouvence* in mammals, we assessed expression by RT-PCR in various mouse and human tissues. The mouse snoRNA-15 was detected in the brain, but not in ovaries, kidney and intestine, while the mouse snoRNA-18 was detected in the intestine, brain, ovaries, but not in the kidney (Figure S7). In human, the unique putative *jou* snoRNA homologue was detected in the intestine, the brain, but weakly in ovaries and kidney.

### RNA-Seq analysis reveals that several genes are deregulated in deletion

To understand *jou* function in longevity we performed a transcriptomic analysis (RNA-Sequencing) of the gut tissue in Wild-Type and jou-deleted flies. Based on a standard stringency of 2-fold changes, RNA-Seq reveals that 376 genes were up-regulated, while 352 genes were down-regulated (Figure 7 and Table S3 and S4). A Gene Ontology (GO) analysis [54] revealed that the majority of the deregulated genes have a catalytic activity (Figure 7C and 7D), and could be categorised in cellular and metabolic processes (Figure 7C,D and Figure S5). We selected some of the most up- or down-regulated genes to perform RT-qPCR on gut tissue. Figure 8 shows that GstE5 (Glutathione S transferase E5), Gba1a (Glucocerebrosidase 1a), LysB (Lysozyme B), and ninaD (neither inactivation nor afterpotential D) are strongly up-regulated in deletion line, whereas CG6296 (lipase and phosphatidylcholine 1-acylhydrolase predicted activities), and Cyp4p2 (Cytochrome P450-4p2) are strongly down-regulated. The regulation of these genes by jou was confirmed by the analysis of the rescued genotypes. For GstE5 gene, the level of RNA is rescued in genomic-rescued line (Del+rescue-1) as well as in Myo-Gal4 expressing *jou* in deletion background (Del,Myo1A>UAS-jou) specifically in enterocytes. For Gba1a and LysB, the rescue is quite similar to GstE5. However, for the ninaD, the regulation seems to be more complex, since the level of RNA is inverted (importantly increased) in genomic-rescued line (Del,rescue-1) and not rescued in Del,Myo1A>UAS-jou line, suggesting that for this gene the fine regulation of the RNA level is more complex and might involve other factors. For the two selected down-regulated genes (CG6296 and Cyp4p2), similar to ninaD, the regulation seems also to be complex since none of them are rescued by the genomic-rescue line (Del,rescue-1) neither by the Myo1A-Gal4 (Del,Myo1A>UAS-jou), although it is partially rescued (statistically different) in Del,rescue-1 and Del,Myo1A>UAS-jou for CG6296. These results suggest that the longevity function of the snoRNA-jou relies on its ability to regulate gene expression in the enterocytes.

**Figure 7.**
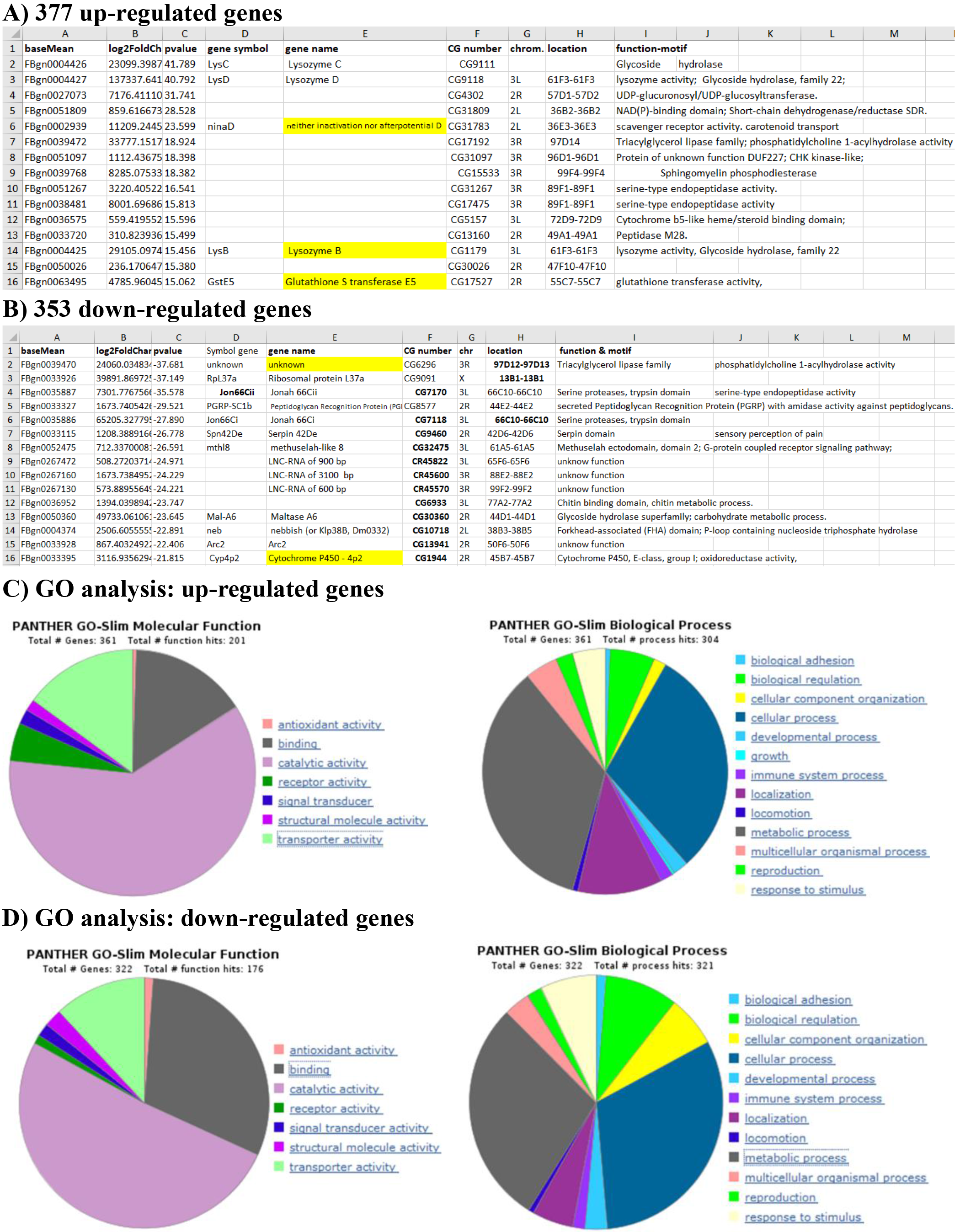
RNA-Seq analysis reveals that hundreds of genes are deregulated in deletion. A) RNA-Seq analysis performed on total-RNA from the gut between the Wilt-Type CS and the deletion reveals that 376 genes are upregulated (A), while 352 are downregulated (B), in deletion compared to Control-CS (Table-S3 and S4 for the complete list of genes). Highlighted in yellow are the selected genes use for validation by RT-qPCR. C&D): Gene Ontology (GO) analysis of the deregulated genes performed with PANTHER classification system [54]. In brief, molecular function selection reveals that the main genes could be classified in catalytic activity, and so both for the up- or down-regulated genes. Selecting for the biological process, the majority of the genes could be grouped into two categories: the cellular and the metabolic processes.

**Figure 8.**
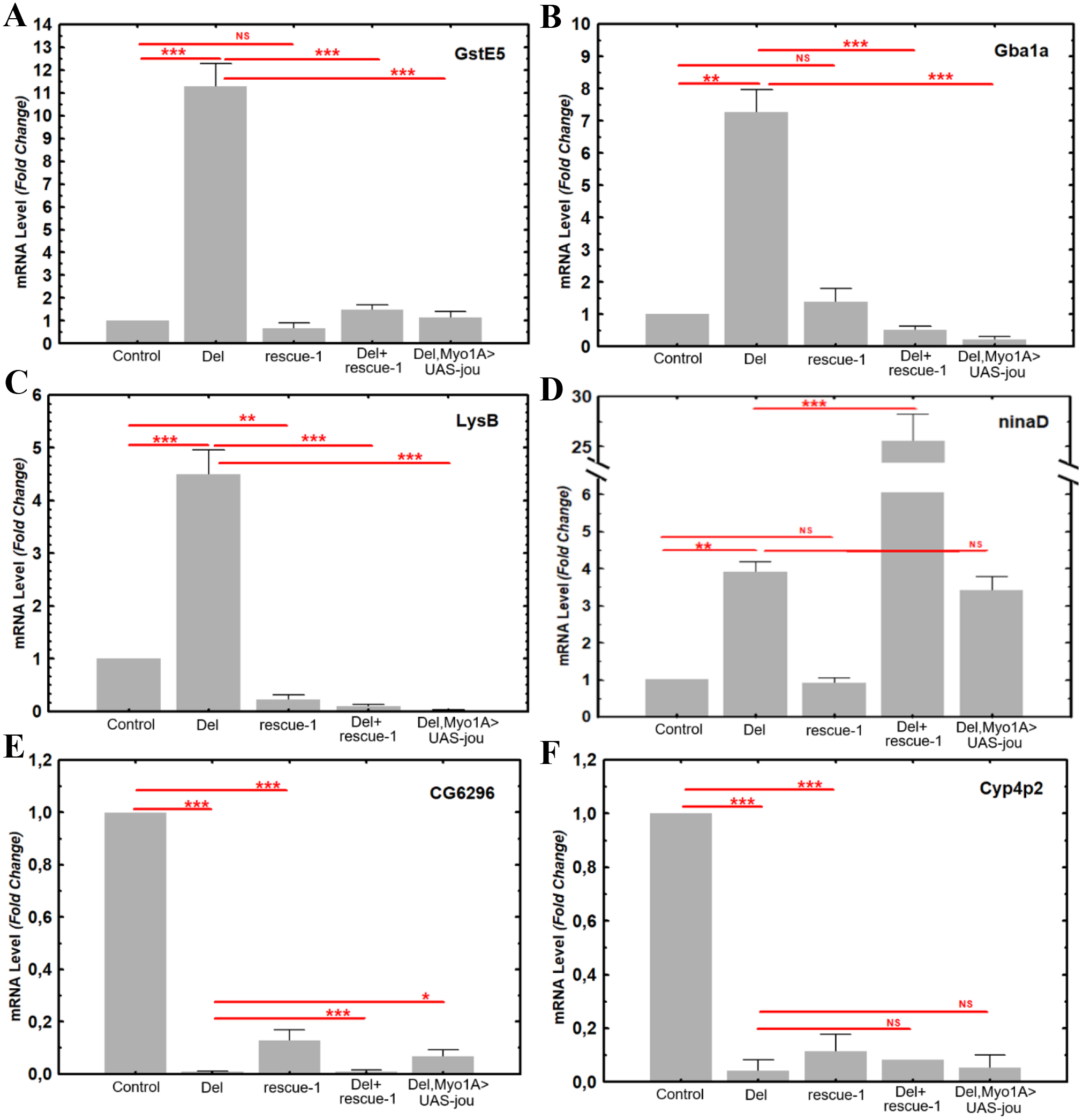
RT-qPCR confirms that genes are deregulated in deletion. RT-qPCR (Sybgreen) results of the quantification of the GstE5, Gba1a, LysB, ninaD, CG6296, and Cyp4p2 genes in Control (CS), Del (deletion F4), rescue-1, Del+rescue-1, and Del,Myo1A>UAS-jou in deletion genetic background. In deletion, the mRNA level is increased for GstE5, Gba1a, LysB, and ninaD genes, while in contrary it is decreased for CG6296 and Cyp4p2. For the genes GstE5, Gba1a, and LysB, the mRNA level is restored in genomic-rescued transgenic flies Del+rescue-1 (even it is more than restored in LysB), while inversely for ninaD, the mRNA level is more increased than in deletion. The targeted expression of *jou* specifically in enterocytes (Del,Myo1A>UAS-jou) in deletion is sufficient to restore the level of mRNA for GstE5, Gba1a and LysB, but again here, not for ninaD. For the down-regulated genes (CG6296 and Cyp4p2), none of the transgenic construction (Del+rescue-1 and Del,Myo1A>UAS-jou) rescues the level of mRNA (although some partial rescued is detectable with Del,Myo1A>UAS-jou for CG6296). Statistics: compared to control and/or to deletion (p-values) (* p<0,05; ** p<0,005; *** p<0,0005).

### The snoRNA *jouvence* modulates the resistance to various stresses

Several stress-response genes have been reported to influence lifespan [5,8,16,17]. Moreover, several of the deregulated genes of the deletion revealed by the RNA-Seq analysis are also known to be involved in various stress resistances (e.g.: Glutathione and Cytochrome P450 family genes) [55–57]. Then, we wonder whether *jouvence* could affect resistance to stress. To elucidate *jou*’s role in stress response, we tested the resistance of transgenic lines (Del+rescue-1 and rescue-1) to desiccation and starvation. During the desiccation test, young flies were placed at 37°C without water, and their survival time was quantified (Figure 6A1). The median survival (50% of dead flies) of control and deletion flies was ~5.5 and ~4.5 hours respectively, showing that the deletion were less robust. However, rescue-1 flies were more resistant, whereas Del+rescue-1 had similar lifespan as control, indicating that the genomic transgene rescued the resistance to stress. Since transgenic flies lived longer than controls, we also performed the desiccation test in aged (40-days-old) flies (Figure 6A2). Similar results were obtained for old flies.

Next, the starvation assay revealed that the median lifespan of control and deletion was ~42 hours and ~50 hours (30% increase), respectively (Figure 6B1). Rescue-1 flies were similar to control, while Del+rescue-1 had a higher lifespan than control, similar to deletion flies. In old flies, both control and deletion were less robust compared to transgenic overexpression (rescue-1) and Del+rescue-1 line (Figure 6B2). These results showed that *jou* deletion are less robust in desiccation but more robust in starvation in young flies, suggesting that *jou’s* protective role depends on the stress test. However, when jou expression was restored in enterocytes under the control of Myo1A-Gal4, resistance to both desiccation (Figure 6C1 and C2) and starvation (Figure 6D1 and D2), was increased, supporting jou function in stress resistance. Together, these results indicate that *jou* is protective against deleterious effects of stress.

## DISCUSSION

Here, we report the identification and the molecular characterization of a novel snoRNA (snoRNA:Ψ28S-1153) that we have named *jouvence* (*jou*), whose deletion decreases lifespan, while inversely, its overexpression increases it. Several independent genetic experiments support this conclusion. First, three independent genomic-rescued transgenic lines (Del+rescue-1; Del+rescue-2; and Del+rescue-3) rescue the longevity in deletion line. Secondly, since ISH revealed that *jou* was expressed in the enterocytes of the epithelium of the gut, targeted expression of *jou* in these specific cells was sufficient to rescue the longevity in the deletion, and even more precisely, only in adulthood (through the use of the Gene-Switch system). Transcriptomic analysis has revealed that several genes are deregulated, among them, genes involved in stress resistance. Thus, while *jou* deleted flies were more sensitive to stress, the rescue of *jou* in deletion increases the resistance to these stress, demonstrating that the presence *jou* in the gut is sufficient to both increase lifespan and protect against stress.

However, while our study was ongoing, a general transcriptomic analysis [41] has revealed two other putative snoRNAs, localized just upstream to the snoRNA:Ψ28S-1153 (*jouvence*). Since the DNA-fragment of 1727 bp used to build the genomic-rescue (rescue-1, 2, & 3) also contains these two putative snoRNAs, thus a putative effect of these two snoRNAs could not be excluded. To verify that the longevity effect was not due to these two putative snoRNAs, we have targeted the expression of the each of them (snoRNA-2 and snoRNA-3 respectively) in the enterocytes. The expression of the each of these snoRNA individually (in the deletion background) did not rescued or lead to an increase of longevity compared to the their co-isogenic control lines, confirming that the longevity effect is due to the snoRNA *jouvence.*

Our findings indicate that *jou* acts in the fly gut epithelium to promote longevity. Interestingly, in *Drosophila*, a relationship between lifespan and the gut was previously reported [58]. In old Wild-Type flies, ISC over-proliferation causes mis-differentiation of ISC daughter cells, resulting in intestinal dysplasia: a phenomenon called “hyperproliferative diseases”. Preserving hyper-proliferation in the gut epithelium extends lifespan [58]. In line with this, two main hypotheses associating *jou* with longevity could be raised. First, *jou* could maintain a better gut homeostasis by increasing and/or optimising gut epithelium regeneration throughout a fly’s life, especially in aged flies. This might improve or optimise nutrient absorption, and enhance metabolism and homeostasis of the organism [58,59]. Second, perhaps a more persistent expression (or an overexpression) of *jou* in enterocytes increase and/or optimise the absorption of nutrient in the gut, all along the life. Indeed, enteric neurons and systemic signals, coupling nutritional status with intestinal homeostasis have been reported [60].

Many longevity genes have been shown to modify the resistance to stress tests [5,8,17]. Interestingly, loss of *jou* function decreases resistance to desiccation, while inversely, the rescue or overexpression of *jou* increased resistance to this stress test. However, in response to starvation, the *jou*-dependent resistance was found to be altered depending on the age of the fly (in deletion: it is increased in young flies, but decreased in old flies compared to Control). Such a difference and even opposite effect in resistance to various stress tests has previously been reported [61], suggesting that the effect on the stress resistance depends rather of the stress test itself rather than being a universal rule of the longevity genes. Indeed, each stress test relies on specific physiological and/or metabolic parameters that might be differently solicited for each stress. Dessiccation is mainly an indicator of the water physiology/homeostasis while the starvation is principally an indicator of the energy metabolism (lipids and/or glucose). In addition, since starvation effect of jou deletion changes during the adult lifespan (increased in young but reduced in old flies compared to controls), this could suggest that energy metabolic components are modified or altered during the course of the adult life.

RNA-Seq and RT-qPCR analyses revealed that several genes are either up-or down-regulated in the deletion. Moreover, for some of these genes (GstE5, Gba1a, and LysB), the level of RNA was rescued by the re-expression of *jouvence* in the enterocytes, demonstrating that *jou* is clearly the snoRNA responsible of this deregulation. However, for some other genes (ninaD, CG6296 and Cyp4p2) the deregulation seems to be more complex, either requiring a very precise level of snoRNA-jou, or potentially involving the two other putative snoRNAs. Nevertheless, these results suggest that the snoRNA *jouvence* might be involved in the regulation of the transcription or as already reported, potentially in the chromatin structure [32].

Based on its primary sequence (Figure 6A), *jou* can be classified in the box H/ACA snoRNA [27]. Interestingly in humans, some pathologies have been associated with various snoRNAs. For instance, a mutation in the H/ACA box of the snoRNA of the telomerase (a RNP reverse transcriptase) yields a pleiotropic genetic disease, the congenital dyskeratose, in which patients have shorter telomeres [33–34]. It has also been reported that the snoRNA HBII-52, a human C/D box type of snoRNA, regulates the alternative splicing of the serotonine receptor 2C [62]. Therefore, we speculate that the mammalian *jou* homologue could play important functions in mouse and human respectively, but it remains to be investigated. Nevertheless, the conservation of sequence of and expression of *jou* raise the exciting hypothesis that similar functions may also be conserved in vertebrates including humans.

Thus, undoubtedly, *jou* represents a good candidate to understand the relationship between the gut and aging/longevity. Interestingly, as already reported, the intestinal cells are very similar in *Drosophila* and mammals, both at cellular and molecular levels [48,63], while snoRNAs are well conserved throughout evolution [27]. Although *jouvence* was not yet annotated in the mammalian genome including human, we have shown that *jou* is present in mammals and is expressed in the gut (among other tissues), strongly suggesting that it might be functional. Therefore, we could hypothesize that manipulating the mammalian gut epithelium may also protect against deleterious effects of aging and even increase longevity (including in humans). Thus, *jou* snoRNA could represent a promising new therapeutic candidate to improve healthy aging.

## EXPERIMENTAL PROCEDURES

### Flies

*Drosophila melanogaster* lines were maintained at 24°C, on our standard dried brewer yeast and cornmeal medium (without sugar or molasses), in a 12h light/12h dark cycle. Canton Special (CS) flies were used as Wild-Type control. The deletion (F4) and the genomic rescue transgenic flies (rescue-1,-2,-3, Del+rescue-1; Del+rescue-2; and Del+rescue-3) were outcrossed a minimum of 6 times with CS (Cantonization) to thoroughly homogenize the genetic background. To follow the deletion (F4) during the Cantonisation (and similar for the Berlinisation), PCR has been performed at each generation to re-identify the fly (chromosome) bearing the mutation (genotyping). The Gal4 lines used (Myo1A-Gal4, Mex-Gal4, and the newly generated Mex-GS line), and the generated transgenic lines (UAS-jou, UAS-sno-2, UAS-sno-3) were also backcrossed 6 times with CS (Cantonization). The two Gal4 lines (Myo1A-Gal4 and Mex-Gal4) were then crossed to each of the snoRNA transgenic construct lines to determine their longevity, while in parallel, they were crossed to deletion (F4) to generate the co-isogenic control flies (in such a way, all the tested lines were in heterozygous both for the Gal4 insertion as well as for the UAS-transgene). The lines Myo1A-Gal4 (located on chromosome 2), esg-Gal4 (chromosome 2), were kindly provided by B.A. Edgar, Heidelberg, Germany. The lines Su(H)GBE-Gal4 (chromosome X), and Dl-Gal4 (chromosome 3) were a courtesy of X. Zeng, NCI/Frederick, USA. Mex-Gal4 (chromosome 2) was kindly provided by G.H. Thomas, Pennsylvania State Univ., PA, USA.

### Determination of the deletion (F4) by PCR

Two primers located at positions 66851F and 68140R (see the Supplementary Information for the sequence of all primers) have been used to amplify the genomic DNA region, both in WT-CS and deletion, to reveal the deletion. The amplicon in CS is 1289 pb, while in deletion is 657 pb. In complement, the deletion has been sequenced. The same two primers have been used to follow the deletion during the Cantonisation and the Berlinisation.

### Construction and genesis of the DNA genomic rescued lines

The *jou* genomic transgenic lines were generated by PCR amplification of a 1723 bp genomic DNA fragment, using the forward primer (snoRNAgenomic-F) and the reverse primer (snoRNAgenomic-R) (see Suppl. Information for the sequence primers), both adding a *XbaI* cloning site at the 5’-ends. The amplified fragment was then inserted via the Xba1 restriction site into the pCaSper 4 (which contains no enhancer/promoter sequence). This way, snoRNA expression is dependent on its own genomic regulatory sequences included in the inserted DNA fragment. The transgenic flies were generated by standard transgenic technique (Bestgene, USA), and three lines have been obtained (rescue-1, rescue-2, and rescue-3). Rescue-1 and rescue-2 are inserted on the third chromosome, while the rescue-3 is inserted on the second chromosome).

### Construction and genesis of the Mex-Gene-Switch line

To generate the Mex-Gene-Switch line, we use the forward primer: Mex1-F + a Sfi1 cloning site at the 5’-ends: 5‘-ATAGGCCGGACGGGCCAACGCGAATTCAGACTGAGC-3’, and the reverse primer: Mex1-R + a Sfi1 cloning site: 5‘-TGTGGCCCCAGTGGCCCGTTGCACATGGTGATGACT-3’, to amplify by PCR using the genomic DNA, a fragment of 2264 bp. Then, this DNA fragment containing the regulatory/promoter sequence of the Mex gene, as formerly reported [50,53] has been inserted through the EcoR1 and DraII sites into the pattB-Sf-HS-GALGS (Sfi1 orientated) (kindly provided by H. Tricoire, Paris). The transgenic flies have been generated by BestGenes (USA), in which the plasmid-construct has been inserted in VK02 site located at position 28E7 on the second chromosome. The Mex-GS transgenic line has been re-introduce in the deletion background by standard genetic cross and recombination.

### Construction and genesis of the UAS-snoRNA lines (jou, sno-2, sno-3)

First, the snoRNA *jou* of 148 bp was cloned into the TOPO-TA cloning vector (Invitrogen) using the same primers used for RT-PCR (see Suppl. Information for snoRNA-Fb and snoRNA-Rb). This construct was also used to synthesize probes for *in-situ* hybridization. Subsequently, the snoRNA fragment of 148 bp was subcloned into the pJFRC-MUH vector (Janelia Farm) containing the attB insertion sites allowing a directed insertion. Similarly, for the cloning of the DNA fragment containing the sno-2 or the sno-3, appropriated forward and reverses primers (see Suppl. Information) were used to amplify a DNA fragment of 166pb for the sno-2, and 157 pb for the sno-3, respectively. Each fragment was then cloned directly into the pJFRC-MUH vector. The transgenic flies were generated (BestGenes, USA) in which the pJFRC-MUH-snoRNA construct was inserted into attP2 site located at position 68A4 on the third chromosome for jou, while the sno-2, and the sno-3 were inserted in VK27 site located at position 89E11 on the third chromosome.

### Detection of *jou* expression by RT-PCR in *Drosophila*, mouse and human

For *Drosophila*, total RNA from whole female flies was extracted by standard procedures using Trizol (Invitrogen). This was treated with RQ1 RNase free DNase (Promega) to remove any DNA contamination, and verified by PCR. Following this, RT was performed with MMLV Reverse Transcriptase (Invitrogen) using a random primer hexamer (Promega), followed by a 20-min RNase H (Invitrogen) at 37°C. The PCR was performed using forward and reverse primers (for primer sequences, see Supplementary Information), with Taq DNA Polymerase (Invitrogen), at 51.4°C. Similar procedures were used to amplify the 300 bp control RP49 fragment using the forward and reverse primers respectively. Mouse and human total RNA were commercially purchased at Amsbio: Mouse Brain, whole Total RNA: MR-201, Mouse Ovary Total RNA: MR-406, Mouse Kidney Total RNA: MR-901, Human Brain Total RNA: HR-201, Human Ovary Total RNA: HR-406, Human Kidney Total RNA: HR-901. The human gut RNA was purchased at Biochain: Human adult Normal small Intestine Total RNA: T1234226. All of these total RNA were re-treated by the DNAses to completely remove the DNA, and re-verified by PCR before performing the RT-PCR. Thereafter, RT-PCR have been performed in similar conditions as for *Drosophila*. For the sequence primers, see Supplementary Information.

### Quantitative PCR (qPCR) by Taqman

For RNA extraction of whole flies or dissected gut, similar procedures described for RT-PCR were used. The samples were diluted in a series of 1/5, 1/10, 1/100 to establish the standard curve, then 1/100 was used for the internal control (RP49) and 1/5 for *jou.* The TaqMan Universal Master Mix II, no UNG was used with a specific Taqman probe designed for the snoRNA *jouvence* and a commercial Taqman probe for RP49 (catalog number: 4331182, ThermoFisher™). The amplification was done an a BioRad apparatus (Biorad CFX96) with a program set at: activation 95°C, 10 min, PCR 40 cycles: 95°C, 15s, 60°C, 1 min. Then, the ratio of the snoRNA (dilution 1/5) was reported to the RP49 (dilution 1/100-since the RP49 is more expressed) and then calculated and normalised to CS = 1. Taqman RT-qPCR was performed in duplicate on each of 3 or 5 independent biological replicates, depending of the genotype. All results are presented as means +/− S.E.M. of the biological replicates.

### *In-situ* hybridization (ISH) and histological preparation

For whole flies ISH, we used 5 day-old females. On day-1 (d1), whole flies were fixed with 4% paraformaldehyde (PFA) for 6 hours at 4°C, and then incubated overnight in 25% sucrose solution at 4°C. On d2, individual flies were embedded in 20% carboxymethylcellulose, frozen in liquid nitrogen, and sectioned using a Cryostat (−20°C) at 30 μm thickness. The sections were rehydrated for 15 min in PBS-1X + 0.1% Tween-20, and post-fixed with 4% PFA for 15 min. They were washed twice with PBS-1X + 0.1% Tween-20, and treated with Proteinase-K (10 μg/ml) for 5 min. Proteinase-K was inactivated using glycine (2 mg/ml) and the sections were washed and post-fixed again. For ISH, the fixed sections were washed with hybridization buffer (50% formamide, 5X SSC, 100 μg/ml heparin, 100 μg/ml of salmon sperm DNA, 0.1% Tween-20) diluted with PBS (1/1) followed by pre-hybridization for 1h at 65°C using the same hybridization buffer. Overnight hybridization with the snoRNA antisense probe (or sense as negative control) was performed at 45°C. The sense and anti-sense probes were synthesized using the DIG RNA labelling kit (SP6 or T7) (Roche™). On d3, the sections were washed four times at 45°C: 3/0-3/1-1/1-1/3 with hybridization buffer/PBS, 15 min each wash. Sections were then washed four times with PBS 1X 0.1% tween, 5 min at RT°, and PBS 1X 0.1% tween + 10% goat serum, 1h, followed by the incubation with the primary antibody, anti-DIG-HRP at 1/100 (Perkin Elmer) for 1h, RT. The immuno-reaction was amplified with the tyramide amplification kit (TSA cyanine 3 plus Evaluation kit-Perkin Elmer) for 8 min in the dark; the sections were washed as usual and counter-stained with DAPI (Roche) for 5 min. The preparation was mounted with Mowiol. For double labelling, following tyramide signal amplification, the sections were incubated with goat serum 2% in PBS 1X 0.1% Tween-20, for 2 hours at RT, and incubated with anti-GFP at 1/500 (Roche), overnight. On d4, the slides were washed, and incubated with the secondary antibody, antimouse FITC at 1/500 (Sigma), washed as usual, and mounted with Mowiol. Same protocol has been used for the ISH on dissected gut, except that we use a FITC-labelled tyramide instead of a Cy3-labelled tyramide.

### Longevity and stress resistance tests

#### Longevity

Following amplification, flies were harvested on one day. Female flies were maintained with males in fresh food vials for 4 days. On day 4, females were separated from males, and distributed in a cage holding between 200 to 250 females. The cage (50 mm diameter x 100 mm high) is made of plexiglass. The top is covered by a mesh, while the bottom (floor) is closed by a petri dish containing the food medium. The petri dish was replaced every two days, and the number of dead flies counted. Note: The lifespan determination of each groups of flies and their respectives controls (genotypes) have been performed simultaneously, strictly in parallel. Moreover, all longevity experiments have been repeated 2 or 3 times independently.

#### Starvation test

Female flies were maintained with males in fresh food vials for 4 days, before testing. On day 4, females were separated from males, and distributed in vials containing 20 females each. Each vial contained a filter, and cotton moisturized with 200 μl of water, to keep the filter moist during the test. The flies were kept at 24°C in a humid chamber, to avoid dehydration. Dead flies were counted every 12 hours.

#### Desiccation test

Flies were reared at 25°C, as usual. Then, female flies were maintained with males in fresh food vials for 4 days, at 25°C, before testing. On day 4, females were separated from males, and distributed in empty glass vials containing 20 females each, without any filter paper. The vials were transferred and tested at 37°C, for young flies. However, since 40 day-old flies die rapidly at 37°C precluding a clear differentiation of the various genotypes, the stress test assay was performed at 30°C, a milder stress condition. Dead flies were counted every hour.

### Transcriptomic analysis (RNA-Sequencing)

For RNA extraction, each single hand-dissected gut was immediately soaked in liquid nitrogen and keep at −80°C. 30 guts were dissected for each genotype. RNA extraction was performed with SV Total RNA Isolation kit (Wizard Purification, Promega) following standard procedures provided by the manufacturer. Thereafter, the total RNA was treated with RQ1 RNase free DNase (Promega) to remove any DNA contamination, and verified by PCR. Then, total RNA were checked for integrity on a bioanalyzer (Agilent). PolyA RNA-Seq libraries were constructed using the Truseq mRNA stranded kit (Illumina) according to the manufacturer recommendations. Libraries were sequenced (Single Read 75pb) on an Illumina NextSeq500 instrument, using a NextSeq 500 High Output 75 cycles kit. Demultiplexing has been done (bcl2fastq2 V2.15.0) and adapters removed (Cutadapt1.9.1). Reads have been mapped on the *D. melanogaster* genome (dmel-all-chromosome-r6.13.fasta) with TopHat2. Mapped reads were assigned to features with featureCounts 1.5.0-p2 and differential analysis has been done with DESeq2. Three independent biological replicates have been done for each genotype [Wild-Type CS and deletion (F4)].

### Quantitative and Statistical Analysis

For the longevity and stress resistance survival curves, log-ranks test was performed using the freely available OASIS software (https://sbi.postech.ac.kr/oasis2/) [64]. For the qPCR, data were analysed statistically using analysis of variance (ANOVA) tests with Statistica™ software.

## AUTHOR CONTRIBUTIONS

JRM conceived and designed the experiments. JRM, LM, SS, KA performed Drosophila experiments and analysed data. CC performed bioinformatics search of jou homologues. JRM wrote the manuscript with input from all authors.

## ACKNOWLEDGMENTS

We thank E. Heard for her help with the search of the mammalian orthologue sequences, A. Bardin, A. Molla-Herman, JR Huynh (Institut Curie, Paris) and F. Rouyer (NeuroPSI, Gif-sur-Yvette) for their helpful comments and critical reading of the manuscript, B.A. Edgar (Heidelberg, Germany), X. Zeng (NCI/Frederick, USA), and G.H. Thomas (Penn. State Univ., USA) for the gut-Gal4 lines, H. Tricoire for the pattB-Sf-HS-GALGS (Gene-Switch vector), and M.N. Soler from Imagif CNRS for her assistance with confocal microscopy. The English language of the preliminary version of this manuscript has been edited by ASK Scientific Ltd, Cambridge, UK, and by A. Bardin for the last version. This work was supported by the CNRS France to JR Martin. The authors declare that they have no competing interests.

## SUPPLEMENTARY INFORMATION

Supplementary Information includes Supplementary Experimental Procedures, four Tables and 7 Figures.

---

**Figure 9.**
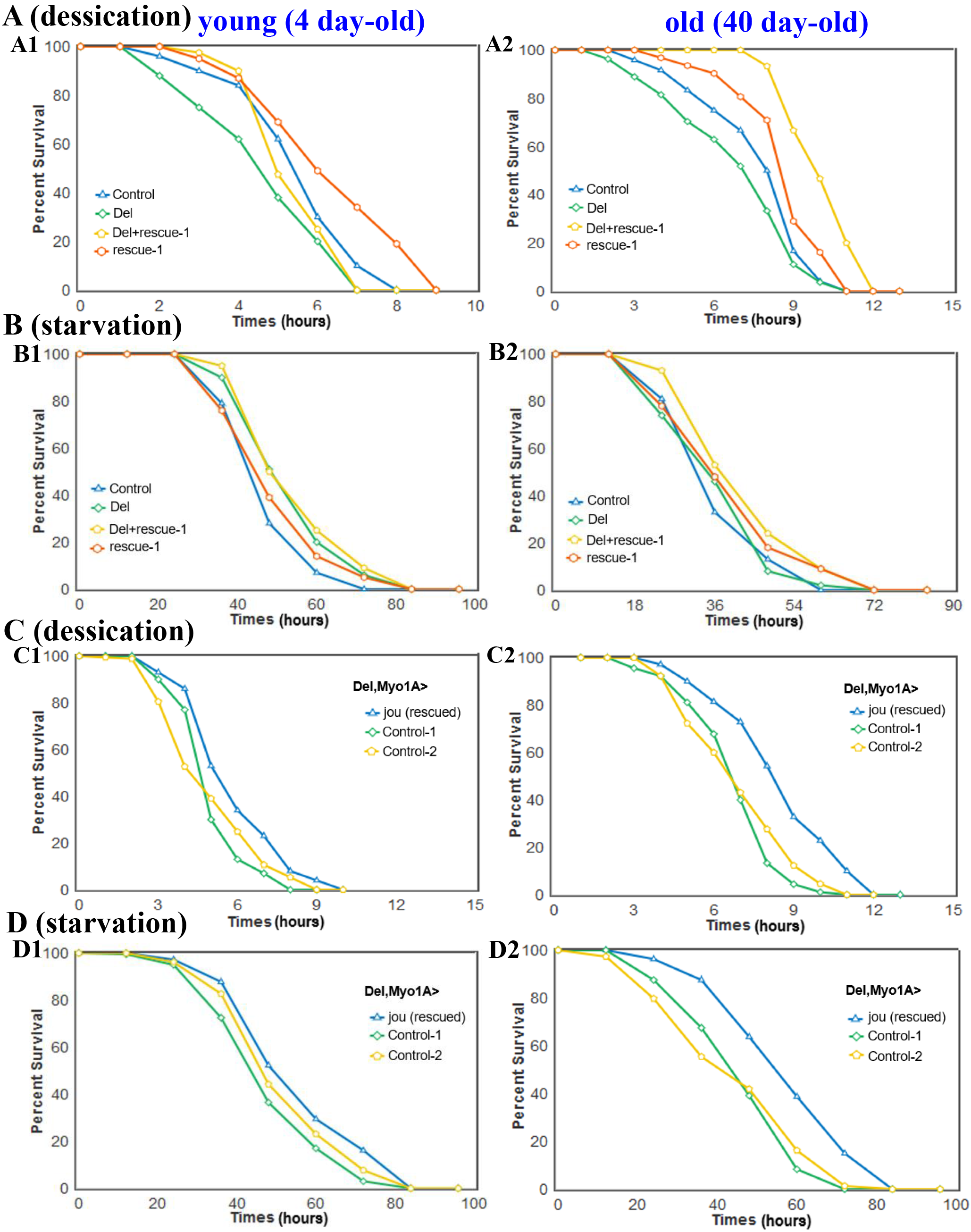
Desiccation and Starvation tests. (A and C) Desiccation and (B and D) Starvation tests in young (A1,B1,C1,D1) and old flies (A2,B2,C2,D2) of genomic-rescued flies (Del+rescue-1) and of enterocytes targeted *jou* through Myo1A-Gal4 line (Del,Myo1A>UAS-*jou*). Both in young and old flies, the targeted expression of the snoRNA-*jou* in the enterocytes increases the resistance to desiccation and starvation, compared to their co-isogenic control flies (Del,Myo1A and UAS-jou/+) (for Statistics: see Table-S2).

## References

1. Fontana, L., Partridge, L. & Longo, V. D. Extending healthy life span--from yeast to humans. Science 328, 321–326 (2010).

2. Niccoli, T. & Partridge, L. Ageing as a risk factor for disease. Curr. Biol. 22, R741–752 (2012).

3. Kennedy, B. K., Austriaco, N. R Jr., Zhang, J. & Guarente, L. Mutation in the silencing gene SIR4 can delay aging in S. cerevisiae. Cell 80, 485–496 (1995).

4. Kaeberlein, M., McVey, M. & Guarente, L. The SIR2/3/4 complex and SIR2 alone promote longevity in Saccharomyces cerevisiae by two different mechanisms. Genes Dev 13, 2570–2580 (1999).

5. Lin, Y. J., Seroude, L., & Benzer, S. Extended life-span and stress resistance in the Drosophila mutant methuselah. Science 282, 943–946 (1998).

6. Clancy, D.J. et al. Extension of Life-Span by Loss of CHICO, a Drosophila Insulin Receptor Substrate Protein. Science 292, 104–106 (2001).

7. Tatar, M. et al. A Mutant Drosophila Insulin Receptor Homolog That Extends Life-Span and Impairs Neuroendocrine Function. Science 292, 107–110 (2001).

8. Broughton, S.J. et al. Longer lifespan, altered metabolism, and stress resistance in Drosophila from ablation of cells making insulin-like ligands. Proc. Natl Acad. Sci. USA 102, 3105–3110 (2005).

9. Bai, H., Kang, P., & Tatar, M. Drosophila insulin-like peptide-6 (dilp6) expression from fat body extends lifespan and represses secretion of Drosophila insulin-like peptide-2 from the brain. Aging Cell 11, 978–985 (2012).

10. Sun, J., & Tower, J. FLP recombinase-mediated induction of Cu/Zn-superoxide dismutase transgene expression can extend the life span of adult Drosophila melanogaster flies. Mol Cell Biol 19, 216–228 (1999).

11. Whitaker, R. et al. Increased expression of Drosophila Sir 2 extends life span in a dose-dependent manner. Aging 5, 682–691 (2013).

12. Rera, M. et al. Modulation of longevity and tissue homeostasis by the Drosophila PGC-1 homolog. Cell Metab 14, 623–634 (2011).

13. Copeland, J.M. et al. Extension of Drosophila life span by RNAi of the mitochondrial respiratory chain. Curr Biol 19, 1591–1598 (2009).

14. Skorupa, D. A., Dervisefendic, A., Zwiener, J. & Pletcher, S. D. Dietary composition specifies consumption, obesity, and lifespan in Drosophila melanogaster. Aging Cell 7, 478–490 (2008).

15. Mair, W., Goymer, P., Pletcher, S. D. & Partridge L. Demography of dietary restriction and death in Drosophila. Science 301, 1731–1733 (2003).

16. Libert, S. et al. (2007). Regulation of Drosophila life span by olfaction and food-derived odors. Science 315, 1133–1137 (2007).

17. Vermeulen, C.J. & Loeschcke, V. Longevity and the stress response in Drosophila. Exp. Gerontol 42, 153–159 (2007).

18. Niccoli, T. et al. Increased Glucose Transport into Neurons Rescues Aβ Toxicity in Drosophila. Curr Biol 26, 2291–2300 (2016).

19. Bolukbasi E et al. Intestinal Fork Head Regulates Nutrient Absorption and Promotes Longevity. Cell Rep 21, 641–653 (2017).

20. Filer, D. et al. RNA polymerase III limits longevity downstream of TORC1. Nature 552, 263–267 (2017).

21. Kenyon, C.J. The genetics of ageing. Nature 464, 504–512 (2010).

22. Chen, Y. F., Wu, C. Y., Kao, C. H. & Tsai, T. F. Longevity and lifespan control in mammals: lessons from the mouse. Ageing Res Rev Suppl 1, S28–S35 (2010).

23. Eacker, S. M., Dawson, T. M. & Dawson, V. L. Understanding microRNAs in neurodegeneration. Nature Rev Neurosci 12, 837–741 (2009).

24. Liu, N. et al. The microRNA miR-34 modulates ageing and neurodegeneration in Drosophila. Nature 482, 519–523 (2012).

25. Bushati, N. & Cohen, S. M. MicroRNAs in neurodegeneration. Curr Opin Neurobiol 18, 292–296 (2008).

26. Kato, M. & Slack, F. J. Ageing and the small, non-coding RNA world. Ageing Res Rev. 12, 429–435 (2013).

27. Kiss, T. Small nucleolar RNAs: an abundant group of noncoding RNAs with diverse cellular functions. Cell 109, 145–148 (2002).

28. Gardner, P.P., Bateman, A. & Poole, A. M. SnoPatrol: how many snoRNA genes are there? J. Biol., 9, 4 (2010).

29. Kiss, T., Fayet-Lebaron, E. & Jády, B. E. Box H/ACA small ribonucleoproteins. Mol Cell 37, 597–606 (2010).

30. Ye, K. H/ACA guide RNAs, proteins and complexes. Curr Opin Struct Biol. 17, 287–292 (2007).

31. Carlile, T.M. et al. Pseudouridine profiling reveals regulated mRNA pseudouridylation in yeast and human cells. Nature 515, 143–146 (2014).

32. McMahon, M., Contreras, A. & Ruggero, D. Small RNAs with big implications: new insights into H/ACA snoRNA function and their role in human disease. Wiley Interdiscip Rev RNA, 6, 173–189 (2015).

33. Mitchell, J.R., Wood, E. & Collins, K. A telomerase component is defective in the human disease dyskeratosis congenita. Nature 402, 551–555 (1999).

34. Vulliamy, T. et al. The RNA component of telomerase is mutated in autosomal dominant dyskeratosis congenita. Nature, 413, 432–435 (2001).

35. Grotewiel, M.S., Martin, I., Bhandari, P. & Cook-Wiens, E. Functional senescence in Drosophila melanogaster. Ageing Res. Rev. 4, 372–397 (2005).

36. Jones, M.A. & Grotewiel, M. Drosophila as a model for age-related impairment in locomotor and other behaviors. Exp. Gerontol. 46, 320–325 (2001).

37. Martin, J.R., Ernst, R. & Heisenberg, M. Temporal Pattern of Locomotor Activity in Drosophila melanogaster. J. Comp. Physiol. A, 184, 73–84 (1999).

38. Brody, T., Stivers, C., Nagle, J. & Odenwald, W. F. Identification of novel Drosophila neural precursor genes using a differential embryonic head cDNA screen. Mech. Dev. 3, 41–59 (2002).

39. Abu-Shumays, R.L. & Fristrom, J. W. IMP-L3, A 20-hydroxyecdysone-responsive gene encodes Drosophila lactate dehydrogenase: structural characterization and developmental studies. Dev. Genet. 20, 11–22 (1997).

40. Huang, Z.P., et al. Genome-wide analyses of two families of snoRNA genes from Drosophila melanogaster, demonstrating the extensive utilization of introns for coding of snoRNAs. RNA, 11, 1303–1316 (2005).

41. Graveley, B. R. et al. The developmental transcriptome of Drosophila melanogaster. Nature 471, 473–479 (2011).

42. Roberts, D. B. Drosophila, A Practical Approach (IRL Press, Oxford University Press, Oxford, U.K). (1998).

43. Brand, A. H., Manoukian, A. S. & Perrimon, N. Ectopic expression in Drosophila. Methods Cell Biol 44, 635–654 (1994).

44. Micchelli, C.A. & Perrimon, N. Evidence that stem cells reside in the adult Drosophila midgut epithelium. Nature 439, 475–479 (2006).

45. Ohlstein, B. & Spradling, A. The adult Drosophila posterior midgut is maintained by pluripotent stem cells. Nature 439, 470–474 (2006).

46. Ohlstein, B. & Spradling, A. Multipotent Drosophila intestinal stem cells specify daughter cell fates by differential notch signaling. Science 315, 988–992 (2007).

47. Jiang, H. & Edgar, B. A. Intestinal stem cells in the adult Drosophila midgut. Exp. Cell Res. 317, 2780–2788 (2011).

48. Takashima, S. et al. Development of the Drosophila entero-endocrine lineage and its specification by the Notch signaling pathway. Dev. Biol. 353, 161–172 (2011).

49. Zeng, X., Chauhan, C. & Hou, S. X. Characterization of midgut stem cell- and enteroblast-specific Gal4 lines in drosophila. Genesis 48, 607–611 (2010).

50. Phillips, M. D. & Thomas, G. H. Brush border spectrin is required for early endosome recycling in Drosophila. J Cell Sci 119, 1361–1370 (2006).

51. Dutta, D. et al. Regional Cell-Specific Transcriptome Mapping Reveals Regulatory Complexity in the Adult Drosophila Midgut. Cell Rep, 12, 346–358 (2015).

52. Roman, G. & Davis, R. L. Conditional expression of UAS-transgenes in the adult eye with a new gene-switch vector system. Genesis 34, 127–131 (2002).

53. Newfeld, S. J., Chartoff, E. H., Graff, J. M., Melton, D. A. & Gelbart, W. M. Mothers against dpp encodes a conserved cytoplasmic protein required in DPP/TGF-beta responsive cells. Development 22, 2099–2108 (1996).

54. Mi, H., Muruganujan, A., Casagrande, J. T. & Thomas, P. D. Large-scale gene function analysis with the PANTHER classification system. Nat Protoc. 8, 1551–1566 (2013).

55. Enayati, A.A., Ranson, H. & Hemingway, J. Insect glutathione transferases and insecticide resistance. Insect Mol Biol. 14, 3–8 (2005).

56. Willoughby, L., Batterham, P. & Daborn, P. J. Piperonyl butoxide induces the expression of cytochrome P450 and glutathione S-transferase genes in Drosophila melanogaster. Pest Manag Sci., 63, 803–808 (2007).

57. Girardot, F.,Monnier, V. & Tricoire, H. Genome wide analysis of common and specific stress responses in adult Drosophila melanogaster. BMC Genomics 5, 74 (2004).

58. Biteau, B. et al. Lifespan Extension by Preserving Proliferative Homeostasis in Drosophila. PLoS Genet. 6, e1001159 (2010).

59. Rera, M., Clark, R. I. & Walker, D. W. Intestinal barrier dysfunction links metabolic and inflammatory markers of aging to death in Drosophila. Proc. Natl Acad. Sci. USA 109, 21528–21533 (2012).

60. Cognigni, P., Bailey, A. P. & Miguel-Aliaga, I. Enteric neurons and systemic signals couple nutritional and reproductive status with intestinal homeostasis. Cell Metab. 13, 92–104 (2011).

61. Cook-Wiens, E. & Grotewiel, M. S. Dissociation between functional senescence and oxidative stress resistance in Drosophila. Exp Gerontol. 37, 1347–1357 (2002).

62. Kishore, S. & Stamm, S. The snoRNA HBII-52 regulates alternative splicing of the serotonin receptor 2C. Science 311, 230–232 (2006).

57. Casali, A. & Batlle, E. Intestinal stem cells in mammals and Drosophila. Cell Stem Cell. 4, 124–127 (2009).

64. Yang, J. S. et al. OASIS: online application for the survival analysis of lifespan assays performed in aging research. PLoS ONE 6, e23525 (2011).

